# Identification of perturbation-responsive regions and genes in comparative spatial transcriptomics atlases

**DOI:** 10.1101/2024.06.13.598641

**Authors:** Alan Yue Yang Teo, Matthieu Gautier, Laurent Brock, Jennifer Y. J. Tsai, Alexandra de Coucy, Achilleas Laskaratos, Nicola Regazzi, Quentin Barraud, Michael V. Sofroniew, Mark A. Anderson, Grégoire Courtine, Jordan W. Squair, Michael A. Skinnider

## Abstract

We introduce Vespucci, a machine-learning method to identify perturbation-responsive regions, genes and gene programs within comparative spatial transcriptomics atlases. We validate Vespucci on simulated and published datasets and show that it outperforms 19 published computational methods for spatial transcriptomics. We apply Vespucci to expose the spatial organization of gene programs activated by therapies that guide repair of the injured spinal cord.

---

Spatial transcriptomics is transforming the molecular interrogation of biological tissues. Existing technologies can reveal the cellular composition of any given tissue, expose the molecular programs expressed across its cytoarchitecture, and resolve the spatial distribution of gene regulation underlying these programs^1–3^. Initial applications of spatial transcriptomics aimed to develop atlases of healthy tissues. The maturation of these technologies is now opening the opportunity to profile tissues across multiple experimental conditions within a single experiment, and consequently, to circumscribe the molecular responses to biological perturbations and diseases within their spatial context^4,5^.

Deriving biological insights from complex spatial transcriptomic datasets requires computational methods that are aligned with the biological rationale of the experiment. For this reason, initial efforts to chart spatial atlases of healthy tissues drove the development of computational methods for tasks such as cellular deconvolution, integration with single-cell transcriptomics, and identification of spatially variable genes^6,7^. The emergence of comparative spatial transcriptomics now necessitates computational approaches to identify regions within a tissue that undergo a response to a biological perturbation, and delineate the individual genes and multi-gene programs that establish this response.

Here, we introduce Vespucci, a machine-learning method to identify perturbation-responsive genes in comparative spatial transcriptomics experiments (**Fig. 1a**). Vespucci first prioritizes the locations within a biological tissue that are un-dergoing the most profound transcriptional responses to a biological perturbation. This prioritization is based on the principle that regions undergoing a transcriptional response to a perturbation become more separable in gene expression space, as compared to less affected regions^8,9^. To quantify this separability, Vespucci trains a random forest classifier to predict the experimental condition from which each spatial barcode was obtained, on the basis of the gene expression profile of this barcode. These predictions are then compared with the true experimental conditions, and spatial locations are prioritized on the basis of the area under the receiver operating characteristic curve (AUC) in cross-validation.

**Fig. 1.**
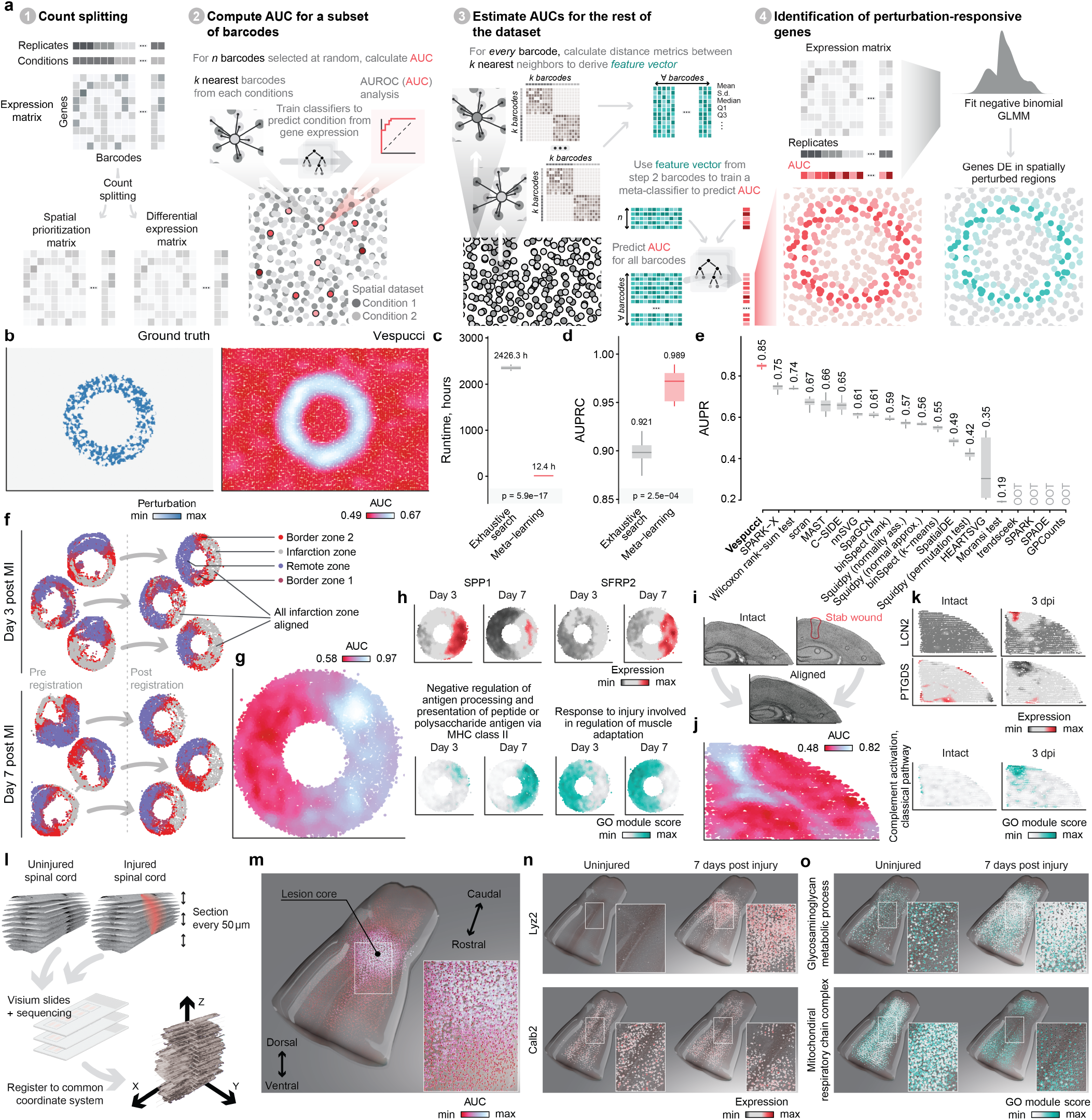
Vespucci identifies perturbation-responsive regions, genes and gene programs in comparative spatial transcriptomics atlases. **a**, Schematic overview of Vespucci. **b**, Benchmarking Vespucci on simulated comparative spatial transcriptomics data. Left, ground-truth pattern of perturbation responsiveness in simulated spatial data. Right, spatial prioritization of transcriptionally perturbed regions in the simulated data, as quantified by the AUC assigned at each spatial coordinate. **c**, Runtime of Vespucci in simulated data, comparing exhaustive calculation of the AUC at each spatial barcode coordinate versus the fast approximation by meta-learning. Inset p-values, two-tailed paired t-test. **d**, Accuracy of spatial prioritization relative to the simulation ground truth, comparing exhaustive calculation of the AUC at each spatial barcode coordinate versus the fast approximation by meta-learning. Inset p-values, two-tailed paired t-test. **e**, Accuracy of spatial prioritization relative to the simulation ground truth, for Vespucci versus 19 baseline methods for SVG identification or spatial DE analysis. **f**, Registration of the myocardial infarction dataset to an annular common coordinate framework separating infarcted and bordering tissues from remote spared tissue. **g**, Spatial prioritization of transcriptionally perturbed regions in the myocardial infarction dataset, as quantified by the AUC assigned at each spatial coordinate. **h**, Expression of selected genes and gene programs (Gene Ontology terms) prioritized by Vespucci in the myocardial infarction dataset. **i**, Registration of the traumatic brain injury dataset to a common coordinate framework. **j**, As in **g**, but for the traumatic brain injury dataset. **k**, As in **h**, but for the traumatic brain injury dataset. **l-n**, As in **i-k**, but for the 3D spinal cord dataset.

Vespucci then reveals the individual genes and multi-gene programs that provide the basis for this spatial prioritization. Given the increasing concern that multiple mechanisms can produce false discoveries when applying differential expression (DE) analysis to single-cell transcriptomics^10,11^ we developed a statistically rigorous approach to identify genes whose expression covaries with the spatial AUC (**Fig. 1a**). To minimize false discoveries, Vespucci tests for DE using a fast negative binomial mixed model^12^ to account for biological and technical variation across replicates, and a count splitting strategy^11^ to avoid overfitting.

To quantify the performance of Vespucci, we simulated comparative spatial transcriptomics data with a well-defined pattern of spatial perturbation. We found that Vespucci accurately recovered the spatial distribution of the simulated perturbation response (**Fig. 1b**). However, achieving this level of accuracy necessitated high-performance computing resources, since analyzing a moderately sized dataset of 10,000 barcodes required more than 2,400 hours of computing time.

We therefore sought to develop a fast approximation that could run on a laptop computer. We devised a meta-learning approach that first computes AUCs for a subset of spatial barcodes, and then predicts the AUCs of the remaining barcodes from a series of distance metrics designed to approximate transcriptional separability. This meta-learning approach accelerated spatial prioritization by more than 200-fold (**Fig.1c**). Unexpectedly, this meta-learning approach also enabled more accurate identification of spatially perturbed regions (**Fig. 1d**).

We next quantified the accuracy with which Vespucci was able to identify the genes that were differentially expressed within the spatially perturbed region. We benchmarked Vespucci against 19 published computational methods developed for spatially variable gene (SVG) identification or spatial DE analysis. Vespucci significantly outperformed all previous methods (p < 10^−5^; **Fig. 1e** and **Extended Data Fig. 1**). This superior performance was maintained across a variety of simulated spatial perturbation pat-terns (**Extended Data Fig. 2**). Moreover, Vespucci maintained control of the false discovery rate when analyzing simulated data without biological differences, whereas several published methods produced thousands of false discoveries (**Extended Data Fig. 3**).

Having validated Vespucci with synthetic data, we next applied Vespucci to reveal perturbation-responsive genes in comparative spatial atlases. We first analyzed a dataset from a mouse model of myocardial infarction^13^, in which spatial transcriptomes were collected at 3 days and 7 days after infarction. We registered all of the sections to a common coordinate system that separated the infarcted zone from bor-dering and remote healthy tissues (**Fig. 1f** and **Extended Data Fig. 4a**). Vespucci recovered the profound transcriptional perturbation occurring within the infarcted region and identified well-studied genes that are known to be involved in the response to myocardial infarction (**Fig. 1g-h** and **Extended Data Fig. 4b**). For instance, Vespucci identified downregulation of *Rpl3l*, a heart-specific ribosomal protein paralog implicated in cardiac-specific translational programs, within the infarcted zone by 7 days^14,15^. Conversely, Vespucci identified upregulation of *Sfrp2*, a gene involved in cardiomyogenesis^16^. Vespucci also implicated less-studied genes in this response, including *Hmox1* and *Hrc* (**Extended Data Fig. 4b-c**).

Beyond the analysis of individual genes, Vespucci can also identify multi-gene modules that are differentially expressed within spatially perturbed regions. In the infarcted heart, Vespucci prioritized gene programs that were upregulated in the infarcted zone or the border region, including programs linked to the adaptive immune response and the response to muscle injury (**Fig. 1h** and **Extended Data Fig. 4b,d**).

We then sought to demonstrate that Vespucci could be applied to biologically and technically heterogeneous spatial datasets. In spatial transcriptomics data from a mouse model of traumatic brain injury^17^, Vespucci prioritized genes related to blood-brain barrier disruption as well as tissue-resident and infiltrating immune cell responses (**Fig. 1i-k** and **Extended Data Fig. 5**). In spatial transcriptomics data from a mouse model of amyotrophic lateral sclerosis^18^, Vespucci prioritized neuronal structural proteins whose downregulation reflected the death of neurons in the gray matter of the spinal cord (**Extended Data Fig. 6**). In spatial transcriptomics data from mice that underwent neurorehabilitation augmented by a spinal cord neuroprosthesis after spinal cord injury^19^, Vespucci prioritized the intermediate laminas of the spinal cord, a finding that reflected the reorganization of specific neuronal populations within these regions, and identified the genes that guide this structural reorganization (**Extended Data Fig. 7**). In an *in situ* RNA sequencing dataset from a mouse model of Alzheimer’s disease^20^, Vespucci prioritized genes whose dysregulation coincided with the deposition of amyloid plaques or hyperphosphorylated tau (**Extended Data Fig. 8**).

We further reasoned that Vespucci could be extended from two to three spatial dimensions in order to scale with the emergence of three-dimensional comparative spatial atlases^4,5^. To demonstrate this possibility, we applied Vespucci to a three-dimensional atlas of the uninjured and injured mouse spinal cord^4^ (**Fig. 1l**). Vespucci recovered the profound transcriptional perturbation within and surrounding the three-dimensional architecture of the lesion site (**Fig. 1m**), and prioritized genes and gene programs associated with calcium signaling, mitochondrial respiration, and glycosaminoglycan metabolism (**Fig. 1n** and **Extended Data Fig. 9**).

Finally, we aimed to demonstrate how Vespucci can support the discovery of new biological mechanisms. To this end, we first deployed Vespucci to study the genes and gene programs activated in response to spinal cord injury in young versus old mice. Whereas young mice can naturally recover the ability to walk after incomplete injury, old mice exhibit permanent paralysis (**Fig. 2a**), and this catastrophic absence of neurological recovery coincides with the failure to reestablish essential neuroprotective barriers between the immune-privileged and extra-neural environments of the injured spinal cord^4^ (**Fig. 2b** and **Extended Data Fig. 10a**).

**Fig. 2.**
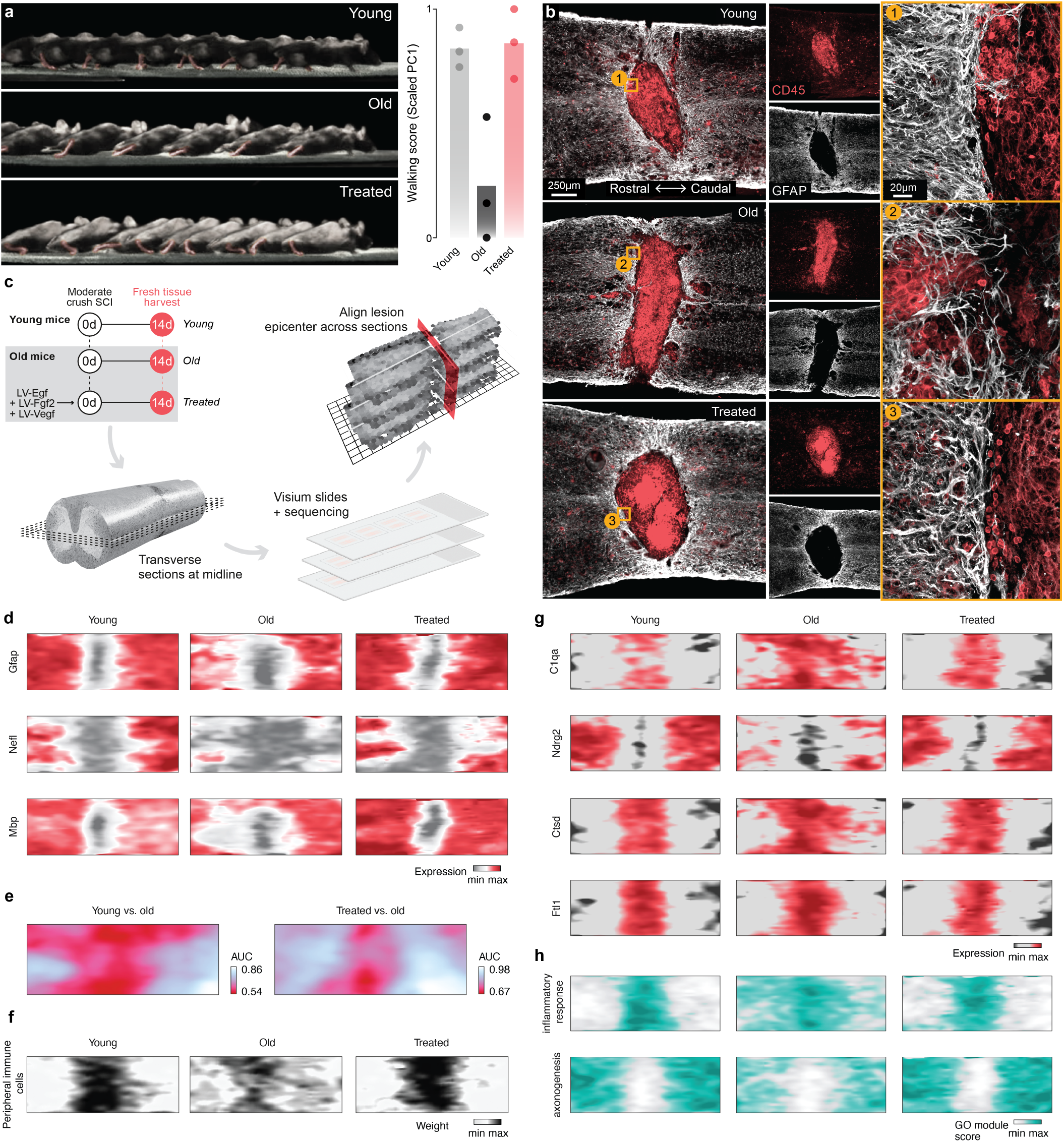
Vespucci reveals genes and gene programs underlying rejuvenation of the aged spinal cord. **a**, Chronophotography of walking in young mice (top), and old mice without (middle) and with (bottom) a gene therapy intervention to promote the formation of the neuroprotective barriers. Bar graph shows the first principal component of kinematic parameters (“walking score”) across groups. **b**, Composite tiled scans of GFAP and CD45 in horizontal sections from representative young, old and treated mice. Insets demonstrate the formation of the astrocyte border, a key component of the neuroprotective barriers that fail to form in old mice but which can be reestablished in treated mice. **c**, Schematic overview of spatial transcriptomics experiments in young, old, and treated mice with spinal cord injury. **d**, Visualization of key histological marker genes on the two-dimensional common coordinate system of the spinal cord. **e**, Spatial prioritization of transcriptionally perturbed regions in injured spinal cords from young versus old mice, left, and old versus treated mice, right. **f**, Cell type deconvolution of comparative spatial transcriptomics data from the injured spinal cord using matched snRNA-seq data, demonstrating the failure to contain peripheral immune invasion in old mice and the reversal of this effect in treated mice. **g**, Expression of selected genes prioritized by Vespucci in comparisons of both young versus old and old versus treated mice. **h**, As in **g**, but showing gene programs (Gene Ontology terms).

To resolve the gene expression programs underlying this failure in the context of the spinal cord cytoarchitecture, we profiled the injured spinal cords of young and old mice by spatial transcriptomics at 2 weeks post-injury. After quality control, all of the sections were registered to a common coordinate system to enable comparative analysis (**Fig. 2c-d** and **Extended Data Fig. 10b-c**).

Vespucci identified the most profound transcriptional perturbation within the spared neural tissues rostral and caudal to the lesion in old mice (**Fig. 2e**), which implicated a failure of old mice to protect tissues adjacent to the injury from peripheral immune invasion, and the ensuing death of spinal cord neurons. Integration of the spatial transcriptomics data with single-nucleus RNA-seq data from young and old mice corroborated this mechanism^21^ (**Fig. 2f**). Vespucci nominated well-studied genes involved in this transcriptional response, highlighting genes associated with dysregulated immune and inflammatory responses (*C1qa, C1qc, Spp1, Tyrobp*) and neuronal death (*Snap25, Ndrg2*) (**Fig. 2g** and **Extended Data Fig. 10d**). These findings were recapitulated when applying Vespucci to expose the gene programs that provided the basis for the spatial prioritization (**Fig. 2h** and **Extended Data Fig. 10d**). Vespucci also highlighted less well-studied aspects of the response to injury, including genes and gene programs associated with iron metabolism (*Ftl1*), proteostasis (*Ctsd, Clu*), and actin formation (*Actb, Tmbs4x*) (**Fig. 2g-h** and **Extended Data Fig. 10d**).

In old mice, the catastrophic failure to establish neuroprotective barriers can be averted by delivering gene therapies that accelerate the formation of these essential neuroprotective barriers by overexpressing growth factors^4^ (**Fig. 2a-b**). We asked whether such a manipulation would reverse the dysregulation of genes and gene programs that were spatially resolved in old mice. To answer this question, we injected lentiviruses into the spinal cords of old mice to overexpress the growth factors *Egf, Fgf2*, and *Vegf* immediately after the injury^4^, and collected spatial transcriptomics sections from the spinal cords of the treated mice (**Fig. 2c-d**). Spatial prioritization of untreated versus treated old mice again prioritized the spared neural tissues adjacent to the lesion, as well as the astrocyte barrier itself (**Fig. 2e**). In keeping with this observation, many of the same genes and gene programs prioritized in old versus young mice were again prioritized when comparing untreated versus treated old mice (**Fig. 2g-h** and **Extended Data Fig. 10e-f**). These findings were consistent with the containment of the lesion and the prevention of neuronal death in treated mice.

Collectively, these results exemplify the power of Vespucci to delineate the landscape of the spatial perturbation-responsive genes and gene programs in comparative spatial atlases, and demonstrate that reestablishing the formation of neuroprotective barriers with gene therapies can reverse age-associated transcriptional dysregulation to repair the damaged spinal cord and restore neurological functions after spinal cord injury.

With spatial transcriptomics now enabling comparative studies of biological perturbations and diseases, Vespucci addresses a key gap in the availability of computational methodologies that are necessary to navigate these complex datasets. A limitation of Vespucci is that the tissue being studied must present a cytoarchitecture that allows sections from different experimental conditions to be registered into a common coordinate system. We envision that Vespucci will help fulfill the potential of comparative spatial atlases to discover new biological principles and accelerate the development of new therapies.

## Methods

### Design and implementation of Vespucci

#### Conceptual framework

The maturation and commercialization of spatial transcriptomics technologies are increasingly allowing investigators to collect comparative spatial atlases that span multiple experimental conditions: for instance, tissues from diseased animals versus healthy controls, or from animals treated with a drug versus untreated controls. We designed Vespucci as a machine-learning method to facilitate the biological interpretation of these comparative spatial atlases. Vespucci first identifies regions of the tissue of interest that are undergoing a transcriptional response to a given perturbation, a process that we refer to as spatial prioritization. Vespucci subsequently identifies the genes or gene programs whose abundance covaries with this transcriptional response to provide the basis for this spatial prioritization (**Supplementary Algorithm 1**).

Vespucci proceeds from a premise that we previously developed and validated in the context of single-cell transcriptomics data: namely, that the cell types undergoing the strongest transcriptional response to a given experimental perturbation are those for which perturbed cells can readily be separated from unperturbed cells by a machine-learning classifier, and that the accuracy with which of this separation can be used as a quantitative mea-sure of perturbation intensity8,^9^. We implemented this concept in a method called Augur, which ranks the cell types identified in a comparative singlecell atlas on the basis of their transcriptional response to a perturbation. In detail, for each cell type annotated in a single-cell atlas, Augur first samples a small number of cells of that type (by default, 20 cells per condition). These cells are partitioned into three folds, and a random forest classifier is trained in threefold cross-validation to predict the condition from which the held-out cells originated (for instance, treated versus untreated). The accuracy of the classifier is quantified by the AUC. The cross-validation procedure is then repeated for many small subsamples (by default, 50 subsamples); we showed that this procedure eliminates the relationship between the AUC and cell type frequency that arises because model performance improves as the size of the training dataset increases. The mean cross-validation AUC across all subsamples then provides a quantitative measure of the intensity of the transcriptional response for cells of that type. The entire procedure is then repeated for each cell type in turn.

Vespucci applies an analogous conceptual framework to identify regions of a comparative spatial transcriptomics dataset that are undergoing a transcriptional response to a perturbation. To this end, we sought to evaluate the transcriptional separability between spatial barcodes from each experimental condition at each (x, y) coordinate in a common coordinate system. We envisioned that this could be achieved by quantifying the separability of spatial barcodes from each condition within small, overlapping tiles, layered across this coordinate system. To this end, Vespucci iterates over every spatial barcode in the dataset and selects its *k* nearest spatial neighbors from each experimental condition, on the basis of the Euclidean distance (*k* is set to 50 by default). Vespucci then divides the selected barcodes into three folds and trains a random forest classifier to predict the experimental condition from which the held-out barcodes were obtained, given the gene expression profiles and experimental conditions of the spatial barcodes in the training folds as input. The accuracy of the predictions is quantified by the AUC, and the process is repeated in three-fold cross-validation. The entire cross-validation procedure is repeated 10 times to converge at a robust estimate of the AUC. This procedure is repeated for each barcode in the dataset, providing a spatial map of the AUC over the coordinate system of the spatial transcriptomics data.

#### Meta-learning

We refer to the strategy described above as “exhaustive search,” because it involves iterating over every spatial barcode and performing repeated cross-validation. For moderately sized spatial datasets, this strategy involves training several hundred thousand classifiers, and consequently requires computational resources that are impractical for routine use. To mitigate this computational demand, we devised a meta-learning strategy to accelerate spatial prioritization. In this approach, the transcriptional separability of a small, random subset of spatial barcodes is first quantified using exhaustive search. Subsequently, a second machine-learning algorithm is trained on the outputs of the classifiers at a subset of spatial barcodes in order to impute the AUCs at the remaining coordinates in the dataset.

Concretely, the meta-learning approach begins by computing a feature vector for every spatial barcode in the dataset. For each barcode *b*, the *k* nearest spatial neighbors in each experimental condition are selected, as described above. This yields two gene expression matrices of dimensions *g* × *k*, one for each experimental condition, where *g* is the number of genes in the dataset. Then, a series of distance metrics are calculated for all pairs of barcodes *b*_*i*,*j*_ to produce a set of *k* × *k* matrices; by default, these metrics include the Pearson correlation, the Spearman correlation, and the codependency index^22,23^. A series of summary statistics (mean, median, first and third quartiles, and standard deviation) are then derived for each distance metric in order to create a feature vector for barcode *b* that summarizes the degree of transcriptional similarity between each pair of spatial barcodes across experimental conditions at that spatial coordinate. These features are then provided as input to a random forest regression model that is trained to predict the AUC calculated by exhaustive search, given the feature vectors and AUCs from a small subset of spatial barcodes as training data. The underlying assumption is that the distance metrics between spatial barcodes from each condition at each coordinate contain sufficient information to estimate the transcriptional separability that is learned directly from the gene expression data in the exhaustive search approach.

To further accelerate the meta-learning approach, we devised a strategy to iteratively expand the number of spatial barcodes in the training set (i.e., those for which AUCs have been calculated by exhaustive search) until convergence has been reached. Concretely, at each step *i*, a set of *n* spatial barcodes (parameter ‘barcodes_per_step’; default, 10) are randomly selected and their AUCs are calculated by exhaustive search. These barcodes are then added to the training set, and the meta-learning regression model is re-trained with the expanded training set. The trained meta-learning model is then applied to impute AUCs for all barcodes, and the vector of AUCs predicted at step i is correlated to the vector of AUCs at the previous step, *i* – 1. This process repeats until the correlation between steps *i* and *i* – 1 exceeds the convergence criterion, or until a user-specified maximum number of barcodes has been subjected to exhaustive search (parameter ‘max_barcodes’; default, 1,000). Convergence is reached when the Pearson correlation coefficient exceeds a threshold (parameter ‘min_cor’; default, 0.8) and the difference between the correlation coefficients at steps *i* and *i* – 1 falls below a user-specified value (parameter ‘epsilon’; default, 5 × 10^−4^) for a given number of steps (parameter ‘steps_tracking’; default, 5).

#### Gene selection

To further improve computational efficiency, Vespucci first filters the input gene expression matrix to remove genes with little variation across spatial barcodes, using a strategy implemented in Augur^8,9^. This strategy entails first fitting a local polynomial regression between the mean and the coefficient of variation^24,25^ and then ranking genes on the basis of their residuals. A fixed proportion of the most highly variable features are retained for each cell type (specified by the ‘var_quantile’ parameter, which defaults to 50% to remove only features that show less-than-expected variation based on their mean abundance). Separately, for each iteration of the repeated cross-validation procedure, a random proportion of features are randomly removed (specified by the ‘feature_perc’ parameter, which also defaults to 50%). In combination, these steps substantially reduce the size of the matrix that must be taken out of a sparse representation for input to the classifier, and therefore also the runtime and memory requirements of the software.

#### Identification of perturbation-responsive genes

Vespucci next seeks to identify the genes that provide the basis for this spatial prioritization. In designing a computational strategy to identify perturbation-responsive genes, we sought to avoid a series of potential pitfalls that have recently been identified in the setting of single-cell transcriptomics. First, recent work has highlighted that, when the same gene expression matrix is used for estimation of a latent variable and then testing individual genes for associations with that latent variable (e.g., unsupervised clustering of cells into cell types followed by DE analysis between cell types), control of the false discovery rate is lost, to the extent that DE genes can be detected between entirely random clusters of cells^11^. We recognized that the same pitfall could arise when using the spatial gene expression matrix both to compute the AUC and subsequently to test for association with the AUC. To avoid this pitfall, we implemented a count splitting strategy^11^, whereby the original count matrix is split into one gene expression matrix *A* for AUC calculation and a second matrix *B* for perturbation-responsive gene identification, as described below. Second, we and others^10,26^ have highlighted the importance of accounting for biological and technical variation across replicates in DE analysis of single-cell transcriptomics data, and demonstrated that failing to account for this variation can lead to thousands of false discoveries. Importantly, we found that both ‘pseudobulk’ approaches and generalized linear mixed models restore control of the false discovery rate. Accordingly, Vespucci employs fast negative binomial generalized linear mixed models, as implemented in the NEBULA package^12^, to test for associations between gene expression and the AUC at each spatial barcode, incorporating replicate (section)-level random effects. This procedure enables the identification of genes that are differentially expressed with respect to the continuous AUC values assigned to each spatial coordinate, i.e., the spatially perturbed regions.

#### Identification of perturbation-responsive gene programs

Vespucci uses an analogous approach to identify gene programs that provide the basis for spatial prioritization. Vespucci first quantifies the average expression of genes associated with each GO term at each spatial barcode using the Seurat function AddModuleScores, which controls for the average expression of randomly selected control features.

For each GO term, the vector of module scores across spatial barcodes is then provided as input to a linear mixed model to identify GO terms that are differentially expressed with the continuous AUC values assigned to each spatial coordinate, incorporating replicate-level random effects. For the analyses presented in the manuscript, GO term annotations were obtained from the Gene Ontology Consortium website, and terms annotated to less than five genes were excluded.

### Benchmarking Vespucci in simulated spatial transcriptomics data

#### Simulations

We benchmarked the performance of Vespucci on simulated spatial datasets, in which the ground truth is known *a priori*. To this end, we adapted code implemented in the Splatter package^27^ to generate simulated spatial transcriptomics data with varying patterns of spatial perturbations. To approximate the conditions that are observed in experimental spatial transcriptomics data, we varied the simulated sequencing depth (by ± 10%) and the magnitude of the perturbation effect size (parameter ‘de.facLoc’, by ± 0.1) across sections. Simulated datasets each contained a total of 10,000 barcodes, distributed evenly between six replicates (three per condition). The simulation parameters were estimated from our large-scale spatial transcriptomic atlas of the injured spinal cord^4^, which was selected owing to the large number of replicates profiled by spatial transcriptomics (total of 36 sections), using the function ‘splatEstimate.’ The simulated spatial patterns, and accompanying parameters, were as follows: for the simulation shown in the main text, the perturbation was simulated within an annular region, with 20% of genes selected for DE (parameter ‘de.prob’) with an effect size (parameter ‘de.facLoc’) of 2. For the second simulation pattern (‘circles’), two partially overlapping regions with distinct perturbation effect sizes were simulated, with de.prob = 0.5 and de.facLoc = 0.5 and 2 for each of the two perturbed regions, respectively. For the third simulation pattern (‘stripes’), three non-overlapping regions with distinct perturbation effect sizes were simulated, with de.prob = 0.5 and de.facLoc = 0.5, 2, and 0.5, respectively. For the fourth simulation pattern (‘flag’), we simulated a dataset with multiple overlapping and non-overlapping regions of distinct effect sizes, with de.prob = 0.5 and de.facLoc = 0.4, 0.8, 1.2, and 2, respectively. The values of de.facScale and de.downProb were held consistent across all simulations at 0.4 and 0.5, respectively. To quantify the performance of Vespucci in correctly recovering the simulated spatial patterns, we evaluated the separation of spatial barcodes within perturbed regions from unperturbed controls using the area under the precision-recall curve. Meta-learning was performed with a maximum of 100 barcodes as the convergence criterion, rather than the default of 1,000. All other Vespucci parameters were set to their default values.

#### False discoveries

To specifically evaluate the control of the false discovery rate, we devised a simulation scenario with strong batch effects but no pattern of biological perturbation. This was achieved by simulating a spatial transcriptomics dataset with 50% of the genes selected for DE, and an effect size of 0.2, but with DE simulated between each of the six sections (rather than between two conditions), and barcodes selected for DE randomly assigned on the coordinate system without any spatial pattern. We then calculated the total number of genes spuriously identified as DE at a 5% false discovery rate.

#### Baseline methods

We carried out an extensive review of the literature to identify existing methods for DE analysis of comparative spatial data or spatially variable gene (SVG) identification, which we refer to collectively as spatial DE methods. Methods that did not provide a hypothesis testing framework (i.e., did not assign a p-value to each gene) were omitted. We identified a total of 19 spatial DE methods implemented in 13 different packages that had been employed by at least one independent publication at the time of our literature review. Methods were considered to be out-of-time (OOT) if the runtime exceeded 72 hours without returning results, and out-of-memory (OOM) if the memory requirements exceeded 128 GB^28^.

#### C-SIDE

C-SIDE^29^ is a statistical framework for cell-type-specific DE analysis of spatial transcriptomics data. DE analysis was performed using the ‘run.CSIDE.single’ function, using the spatial coordinates as a random explanatory variable, assigning all barcodes to a single cell type, and with default parameters.

#### Giotto

Giotto^30^ performs dimensionality reduction on the input spatial transcriptomics data, and then constructs a spatial Delaunay triangulation network to delineate the spatial relationships between each barcode. The data was normalized using the ‘normalizeGiotto’ function, with default parameters. Giotto then provides two separate methods for SVG identification (‘binSpect_kmeans’ and ‘binSpect_rank’), as well as wrapper functions to two other SVG detection methods adapted from the scran31 and MAST32 packages. For binSpect_kmeans, expression values for each gene are binarized with k-means clustering, whereas for binSpect_rank, expression values are binarized using a rank-based threshold. All methods were run with default parameters.

#### Gpcounts

GPcounts^31^ uses Gaussian process regression to model spatial transcriptomics data using a negative binomial likelihood function. Spatial coordinates and the gene expression count matrix were provided as input to the ‘Fit_GPcounts’ function, and hypothesis testing was performed using the ‘One_sample_test’ function. All functions were run with default parameters.

#### HEARTSVG

HEARTSVG^32^ uses a series of autocorrelation-based statistics to identify SVGs and then pools the results of four tests using Stouffer’s method. The method was run with default parameters.

#### nnSVG

nnSVG^33^ fits nearest-neighbor Gaussian process models to normalized gene expression values for each gene, then uses likelihood ratio tests to identify spatially variable genes. The gene expression matrix was normalized using the ‘logNormCounts’ function from scater^34^, and the ‘nnsvg’ function was run with default parameters.

#### Seurat

Seurat^35^ provides two workflows for SVG identification in their 10x Visium vignette. In the first workflow, the ‘FindMarkers’ function is used to test for DE genes (e.g., between conditions or clusters of spatial barcodes) without explicitly incorporating spatial information. For this approach, the gene expression matrix was normalized using the ‘Normalize-Data’ function, and ‘FindMarkers’ was run with default parameters, using the Wilcoxon rank-sum test to identify DE genes between experimental conditions. In the second workflow, the ‘FindSpatiallyVariables’ function is used to identify genes that display spatially variable gene expression. For this approach, the gene expression matrix was normalized using SCTransform^36^, and ‘FindSpatiallyVariableFeatures’ was run with selection.method = ‘moransi’, using Moran’s I test to identify genes with significant autocorrelation.

#### SPADE

SPADE^37^ fits Gaussian process regression models with genespecific Gaussian kernels, and performs hypothesis testing using a crossed likelihood-ratio test. The method was run with default parameters.

#### SpaGCN

SpaGCN^38^ identifies spatial domains using a graph convolutional neural network to integrate gene expression, spatial coordinates, and (optionally) histological data, and then performs DE analysis between domains using the Wilcoxon rank-sum test. The ‘rank_gene_groups’ function was run with default parameters to identify DE genes, using experimental conditions rather than spatial domains to define the comparison of interest.

#### SPARK

SPARK^39^ models spatial count data through generalized linear spatial models and tests a series of null hypotheses based on ten different spatial kernels, then combines p-values across all ten spatial kernels via conversion to Cauchy statistics. Covariates were estimated using the ‘spark.vc’ function and statistical testing was performed using ‘spark.test,’ both with default parameters.

#### SPARK-X

SPARK-X^40^ is an update to SPARK that tests for dependence of gene expression on spatial coordinates using a covariance test framework that incorporates a variety of spatial kernels. The ‘sparkx’ function was run with default parameters.

#### SpatialDE

SpatialDE^41^ uses Gaussian process regression to decompose expression variability into spatial and nonspatial components and performs hypothesis testing using a likelihood ratio test, compared to a null model without spatial covariance. The method was run with default parameters.

#### Squidpy

Squidpy^42^ employs Moran’s I spatial autocorrelation statistics to identify spatially variable genes. P-values can be computed in one of three different ways (based on permutations, based on standard normal approximation from permutations, or under normality assumption). We compared all three approaches, using the ‘spatial_autocorr’ function with n_perms = 100.

#### Trendsceek

Trendsceek^43^ models gene expression data as marked point processes and tests for significant dependency between the spatial distributions of points and gene expression levels at those points through pairwise analyses of points as a function of the distance between them. Hypothesis testing is performed via permutation tests. The implementation from the Giotto package (https://github.com/RubD/Giotto/blob/HEAD/R/spatial_genes.R) was adapted to perform mark estimation with ‘set_marks’ and statistical testing using ‘calc_trendstats,’ both with default parameters.

### Preprocessing and analysis of published spatial datasets

#### Myocardial infarction

Calcagno et al.^13^ subjected adult mice to left anterior descending coronary artery ligation and profiled hearts at various time points, along with sham controls, by spatial transcriptomics (10x Visium). We focused on the comparison of three days versus seven days post-MI because these were the two timepoints at which more than one replicate was profiled (*n* = 3 samples per timepoint). Data was obtained from the GEO (accession GSE214611). The six tissue samples were registered to a common coordinate framework to permit comparison across conditions. This was achieved by first aligning all spatial transcriptomics sections to an annulus using the R image registration package RNiftyReg, and then manually rotating each sample in order to align the lesion sites, which were identified by visual inspection of the histological images, quality control statistics (e.g., number of UMIs), the expression of collagen genes, and cell type deconvolution of the spatial data using matched single-cell and single-nucleus RNA-sequencing data collected by the authors from the same mouse models, which were obtained from the same GEO accession.

#### Traumatic brain injur

Koupourtidou et al.^17^ subjected adult mice to traumatic brain injury, using a stab wound model, and profiled coronal brain sections containing the cortex, white matter, and hippocampal formation from injured mice at three days post-injury and uninjured controls (*n* = 2 per condition). Data was obtained from the GEO (accession GSE226208). The four tissue samples were registered to a common coordinate framework to permit comparison across conditions. This was achieved by first manually rotating each sample in order to align the lesion sites, which were identified by visual inspection of the histological images, quality control statistics (e.g., number of UMIs), and then aligning the injured spatial transcriptomics sections to the uninjured sections using the R image registration package RNiftyReg.

#### ALS

Maniatis et al.^18^ profiled lumbar spinal cord tissue sections from a mouse model of ALS (SOD1-G93A mice) and wild type controls by spatial transcriptomics (GEO accession GSE120374). We focused on the comparison of mice at postnatal day 100 to wild type mice. The sections were registered to a common coordinate framework by the authors in their original study, and the coordinates of each spatial barcode were obtained from the authors directly. A small number of barcodes with coordinates well outside of the spinal cord tissue itself (spatial outliers) were removed manually. In view of the left-right symmetry of the spinal cord, we replaced the coordinate of each barcode on the x-axis with its absolute value in order to mitigate artefacts introduced by the registration procedure, as previously described19.

#### Neuroprosthetic rehabilitation

Kathe et al.^18^ subjected adult mice to spinal cord injury, and a subset of the injured mice were additionally subjected to a number of rehabilitative and/or neuroprosthetic interventions. Lumbar spinal cord sections from these mice were then profiled by spatial transcriptomics (10x Visium). Here, we focused on the comparison of injured mice to those that received both epidural electrical stimulation (EES) of the lumbar spinal cord and four weeks of neurorehabilitative training (EES^REHAB^). Data was obtained from the GEO (accession GSE184369), including the coordinates of each spatial barcode in the common coordinate framework established in the original study.

#### Alzheimer’s disease

Zeng et al.^20^ analyzed brain tissues from TauPS2APP triple transgenic mice, a mouse model of Alzheimer’s disease, at 8 and 13 months of age. Coronal sections were profiled by STARmap PLUS, an *in situ* spatial transcriptomics method that targeted 2,766 genes curated from previous bulk and single-cell RNA-seq studies or based on a known or suspected role in AD. Data were obtained from the Zenodo deposition provided by the authors (doi: 10.5281/zenodo.7332091). The sections were registered to a common coordinate framework by using a subset of the cell types defined by the authors of the original study (CA1, CA2, and DG neurons) as markers for registration. To this end, spatial barcodes were assigned categorical labels based on their assignments to one of these three cell types or any other cell type. Every pair of binarized images was then enumerated and registered to one another using the R image registration package RNiftyReg. The final coordinates were obtained by averaging the registered coordinates from each pair of images.

#### 3D spatial atlas of spinal cord injury

Skinnider et al.^4^ collected serial sections at 50 µm intervals throughout the entire dorsoventral axis of the spinal cords of uninjured mice and mice subjected to crush spinal cord injury at 7 days or 2 months, and profiled the sections by spatial transcriptomics. We focused here on the comparison of 7 days post-injury versus uninjured spinal cords. Data was obtained from the GEO (accession GSE234774), including the coordinates of each spatial barcode in the common coordinate framework established in the original study. Vespucci was extended to threedimensional spatial prioritization by setting the value of the ‘coords_col’ parameter to a vector containing the x, y, and z dimensions.

### Comparative spatial transcriptomics of the injured spinal cord

#### Mouse model

Adult male or female C57BL/6 mice (15-25 g body weight, 8-15 weeks of age) were used for all experiments. Aged mice were purchased from JAX at 60 weeks of age (stock no. 000664). Housing, surgery, behavioral experiments and euthanasia were all performed in compliance with the Swiss Veterinary Law guidelines. Manual bladder voiding and all other animal care was performed twice daily throughout the entire experiment. All procedures and surgeries were approved by the Veterinary Office of the Canton of Geneva (Switzerland; authorizations GE/145/2).

Spinal cord crushes were performed at the level of T10, as previously described^44,45^. Animals were euthanised at 14 days post-injury. Crush spinal cord injuries were introduced at the level of T10/T11 after laminectomy of a single vertebra by using No. 5 Dumont forceps (Fine Science Tools) with a spacer so that when closed a 0.5 mm space remained, and with a tip width of 0.5 mm to completely compress the entire spinal cord laterally from both sides for 5 s.

#### Biological repair intervention

Viruses used in this study were produced at the EPFL Bertarelli Foundation Platform in Gene Therapy and included SIN-cPPT-PGK-FGF2-WPRE,SIN-cPPT-PGK-EGF-WPRE SIN-cPPT-PGK-VEGF-WPRE, SIN-cPPT-GFAP-GDNF-WPRE. General surgical procedures have been previously described in detail^44–46^. Surgeries were performed at EPFL under aseptic conditions and under 1-2% isoflurane in 0.5-1 L/min flow of oxygen as general anesthesia, using an operating microscope (Zeiss) and rodent stereotaxic apparatus (David Kopf) as previously described^45,46^. LV injections were made the same day as the SCI, and were targeted over the intended spinal cord segment to be injured. LVs were injected into four sites (two sets of bilateral injections, 0.30 µL/injection [all vectors diluted to 600 µg P24/mL in sterile saline]) 0.6 mm below the surface at 0.15 µL per minute using glass micropipettes connected via high-pressure tubing (Kopf) to 10 µL syringes under the control of a microinfusion pump. After surgeries, mice were allowed to wake up in an incubator. Analgesia, buprenorphine (Essex Chemie AG, Switzerland, 0.01-0.05 mg/kg s.c.) or carprofen (5 mg/kg s.c.), was given twice daily for 2-3 days after surgery. Animals were randomly assigned numbers and thereafter were evaluated blind to experimental conditions.

#### Behavioural assessments

Kinematics data from young, old, and treated mice is reproduced from ref.^4^. Behavioral procedures have been previously described in detail^46–48^. Briefly, during overground walking, bilateral leg kinematics were captured with twelve infrared cameras (Vicon Motion Systems) that tracked reflective markers attached to the crest, hip, knee, ankle joints, and distal toes. The limbs were modeled as an interconnected chain of segments and a total of 80 gait parameters were calculated from the recordings. To evaluate differences between experimental conditions, as well as to identify the most relevant parameters to account for these differences, we implemented a multistep multifactorial analysis based on principal component analysis, as previously described in detail^46,47,49^, and coupled to automated, markless tracking software^50^. Reconstructed kinematic data was processed with custom R scripts to compute gait parameters. For each experiment, a principal component analysis was performed by computing the covariance matrix *A* of the ensemble of parameters over the gait cycle, after subtraction of their respective mean values. The principal components were computed from eigenvalues *λ*_*j*_ and eigenvectors *U*_*j*_ of A. The principal components were ordered according to the amount of data variance accounted for by each component. The coordinate of each gait cycle on the first principal component, i.e., the component vector explaining the greatest amount of variance across the gait parameters, was thereafter referred to as the walking score. These scores were subsequently normalized for each experiment. Individual parameters were then selected to be compared between groups based on their correlation to the first principal component.

#### Perfusions

Mice were perfused at the end of the experiments. Mice were deeply anesthetized by an intraperitoneal injection of 0.2 mL sodium pentobarbital (50 mg/mL). Mice were transcardially perfused with PBS followed by 4% paraformaldehyde in PBS. Tissue was removed and post-fixed overnight in 4% paraformaldehyde before being transferred to PBS or cryoprotected in 30% sucrose in PBS.

#### Immunohistochemistry

Immunohistochemistry was performed as previously described^44–46^. Perfused post-mortem tissue was cryoprotected in 30% sucrose in PBS for 48 h before being embedded in cryomatrix (Tissue Tek O.C.T., Sakura Finetek Europe B.V.) and freezing. 30 µm thick transverse or horizontal sections of the spinal cord were cut on a cryostat (Leica), immediately mounted on glass slides and dried or in free floating wells containing PBS plus 0.03% sodium azide. Primary antibodies were: rabbit anti-GFAP (1:1000; Dako) and rat anti-CD45 (1:100, BD Biosciences). Fluorescent secondary antibodies were conjugated to Alexa 555 (red) or Alexa 647 (far red) (ThermoFisher Scientific). The nuclear stain was 4’,6’-diamidino-2-phenylindole dihydrochloride (DAPI; 2 ng/mL; Molecular Probes). Sections were imaged digitally using a slide scanner (Olympus VS-120 Slide scanner) or confocal microscope (Zeiss LSM880 + Airy fast module with ZEN 2 Black software). Images were digitally processed using ImageJ (ImageJ NIH) software or Imaris (Bitplane, version 9.0.0).

#### Chronophotography

Chronophotography was used to generate a representative series of still pictures arranged in a single photograph to illustrate the walking of mice. Videos at 25 fps or photographs at 15 fps were recorded while mice were performing tasks such as quadrupedal walking on the runway. Images from these recordings were chosen to best illustrate the different consecutive phases of walking of the hindlimbs, i.e. stance phases and swing phases. The frequency of chosen pictures varied due to the varying velocity of the mice. The series of pictures were assembled in Photoshop while blending out non-essential details.

#### Spatial transcriptomics library preparation

For each experimental condition, we prepared sections from the lesion epicenter of four independent biological replicates. Spinal cords were embedded in OCT and cryosections were generated at 10 µm at –20°C. Sections were immediately placed on chilled Visium Tissue Optimization Slides (catalog no. 1000193, 10x Genomics) or Visium Spatial Gene Expression Slides (catalog no. 1000184, 10x Genomics). Tissue sections were then fixed in chilled methanol and stained according to the Visium Spatial Gene Expression User Guide (catalog no. CG000239 Rev A, 10x Genomics) or Visium Spatial Tissue Optimization User Guide (catalog no. CG000238 Rev A, 10x Genomics). For gene expression samples, tissue was permeabilized for 12 min. Brightfield histology images were taken using a 10× objective on a slide scanner (Olympus VS-120 Slide scanner). For tissue optimization experiments, fluorescent images were taken with a TRITC filter using a 10× objective and 400 ms exposure time. Libraries were prepared according to the Visium Spatial Gene Expression User Guide. Samples were sequenced on our NovaSeq 6000 (EPFL Gene Expression Core Facility) with a target of 50,000 reads per spot, according to 10x Genomics guidelines.

#### Read alignment and quality control

Read alignment was performed using SpaceRanger (10x Genomics, version 2.1.1) to obtain a UMI count matrix. Quality control (QC) metrics, including the number of UMIs, number of genes detected, and proportion of reads aligned to mitochondrial genes, were then calculated for each spatial barcode using Seurat^35^. Low-quality sections were identified as those with unusually low numbers of UMIs or genes expressed, or QC metrics that differed markedly from the remainder of the sections in the dataset. Removal of low-quality sections afforded a final dataset comprising eight sections (three from young mice, three from old mice, and two from treated old mice).

#### Registration to a common coordinate system

We aligned all of the spatial transcriptomics sections into a common coordinate system using a custom image analysis pipeline. This pipeline involved preprocessing, registration, and merging of the histological images. Image preprocessing was conducted in Fiji, while registration procedures were executed in R using the ‘imager’ package. A custom macro in Fiji was used to segment the histological sections and their spatial barcodes from the background. The segmented sections were then aligned with ‘imager’. Manual image registration was performed based on the tissue structure, which was identified by inspection of (i) histological images, (ii) quality control statistics (e.g., mitochondrial count percentages), and (iii) marker genes for the major cell types of the spinal cord (e.g., *Nefl*).

#### Cell type deconvolution

RCTD^21^ was used to deconvolve spatial barcodes into a mixture of one or more cell types, while accounting for technical differences between single-nucleus and spatial transcriptomes, using published snRNA-seq data from the spinal cords of old and young mice at seven days after crush SCI^4^. A single cell type was assigned to each spatial barcode by taking the maximum deconvolution weight assigned by RCTD for that barcode, and the resulting patterns of cell type abundance were smoothed by 2D locally weighted regression as described by the authors of RCTD^21^.

### Visualization

Spatial AUCs, gene expression values, and GO module scores were smoothed prior to visualization in the two-dimensional spatial transcriptomics datasets using locally weighted regression, as implemented in the RCTD package^21^. If outliers were still present, the smoothed values were winsorized at the first and 99th percentiles to improve visualization. To visualize the three-dimensional spatial dataset, we first exported spatial AUCs, gene expression values, and GO module scores to CSV files, and then imported these into Blender using the spreadsheet importer tool at https://github.com/simonbroggi/blender_spreadsheet_import. The data was then visualized using geometry nodes and point clouds, with colors assigned using custom materials that took quantitative values as input to a color ramp node. Throughout the paper, boxplots show the median (horizontal line), interquartile range (hinges) and smallest and largest values no more than 1.5 times the interquartile range (whiskers).

## Code availability

Vespucci is available from GitHub at https://github.com/neurorestore/Vespucci.

## Acknowledgements

This work was supported by the Swiss National Science Foundation (grant nos. 310030_192558 to G.C., 320030_228288 to M.A.A. and PZ00P3_208988 to J.W.S.); Wings for Life (to M.A.S., M.A.A. and M.V.S.); Friedrich Flick Förderungsstiftung through Wings for Life (to M.A.A. and G.C.); Wyss Center for Bio and Neuroengineering (to M.A.A. and G.C); endParalysis Foundation (to M.A.A.); and the ALARME Foundation (to M.A.A. and G.C.).

## Author contributions

A.Y.Y.T., G.C., J.W.S., and M.A.S. conceived and designed experiments. A.Y.Y.T., M.G., L.B., J.Y.J.T., A.d.C., A.L., N.R., Q.B., M.A.A., J.W.S., and M.A.S. conducted experiments. A.Y.Y.T., M.G., L.B., J.Y.J.T., Q.B., M.V.S., M.A.A., J.W.S., and M.A.S. analyzed the data. A.Y.Y.T., G.C., J.W.S., and M.A.S. wrote the manuscript. All authors contributed to the editing of the manuscript.

---

**Extended Data Fig. 1.**
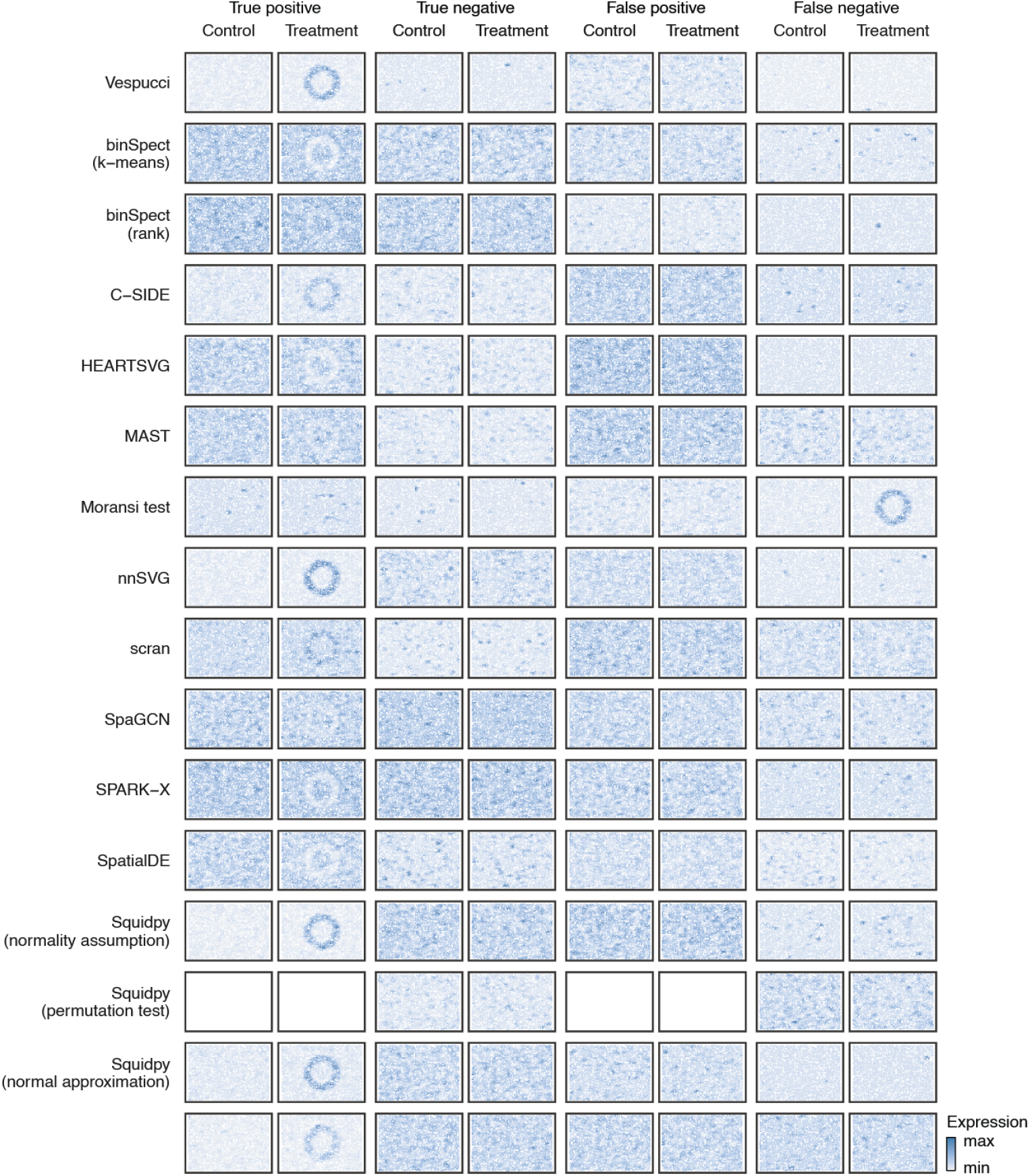
Examples of genes prioritized by Vespucci versus 19 baseline methods in simulated comparative spatial data. A single representative gene from each of four categories (true positive, true negative, false positive, false negative) was selected at random from the output of each computational method. Blank spaces indicate methods that did not identify any genes from a given category.

**Extended Data Fig. 2.**
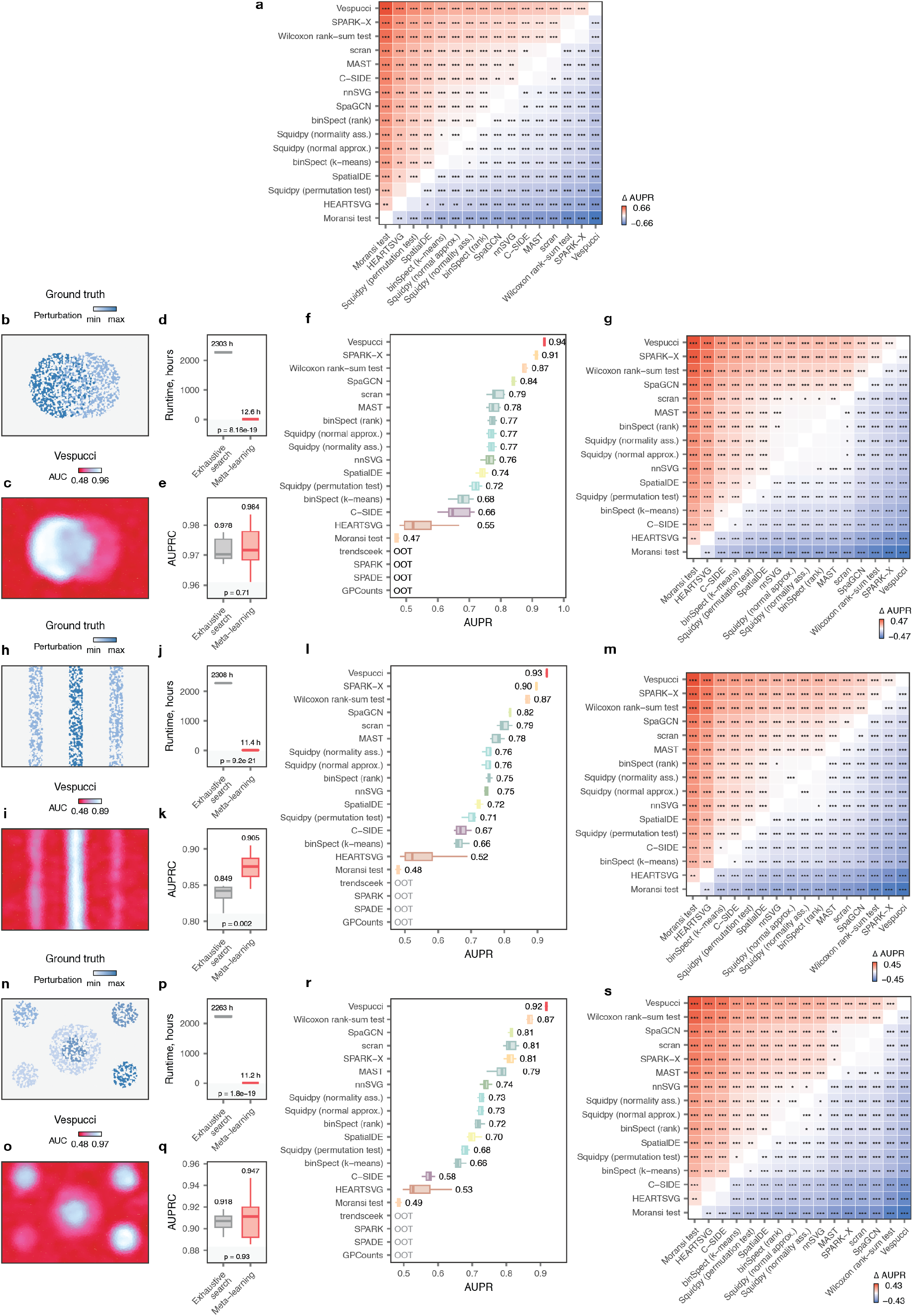
Evaluation of Vespucci in alternative simulated spatial perturbations. **a**, Accuracy of spatial prioritization relative to the simulation ground truth for the simulated spatial perturbation shown in **Fig. 1**, for Vespucci versus 19 baseline methods for SVG identification or spatial DE analysis, as quantified by the area under the precision-recall curve (AUPR). Inset p-values, two-tailed paired t-test. **b**, Ground-truth pattern of perturbation responsiveness in a second simulated spatial dataset with two partially overlapping perturbations of unequal magnitudes (“circles”). **c**, Spatial prioritization of transcriptionally perturbed regions in the simulated data, as quantified by the AUC assigned at each spatial coordinate. **d**, Runtime of Vespucci in the simulated data, comparing exhaustive calculation of the AUC at each spatial barcode coordinate versus fast approximation by meta-learning. **e**, Accuracy of spatial prioritization relative to the simulation ground truth, comparing exhaustive calculation of the AUC at each spatial barcode coordinate versus the fast approximation by meta-learning. Inset p-values, two-tailed paired t-test. **f**, Accuracy of spatial prioritization relative to the simulation ground truth, for Vespucci versus 19 baseline methods for SVG identification or spatial DE analysis, as quantified by the area under the precision-recall curve (AUPR). Inset p-values, two-tailed paired t-test. **g**, Heatmap comparing the mean differences in AUPR for each pair of computational methods. *, p < 0.05; **, p < 0.01; ***, p < 0.001, two-tailed paired t-test. **h-m**, As in **b-g**, but for a third simulated spatial dataset with three non-overlapping perturbations of unequal magnitudes (“stripes”). **n-s**, As in **b-g**, but for a fourth simulated spatial dataset with multiple perturbations of unequal magnitudes, some partially overlapping (“flag”).

**Extended Data Fig. 3.**
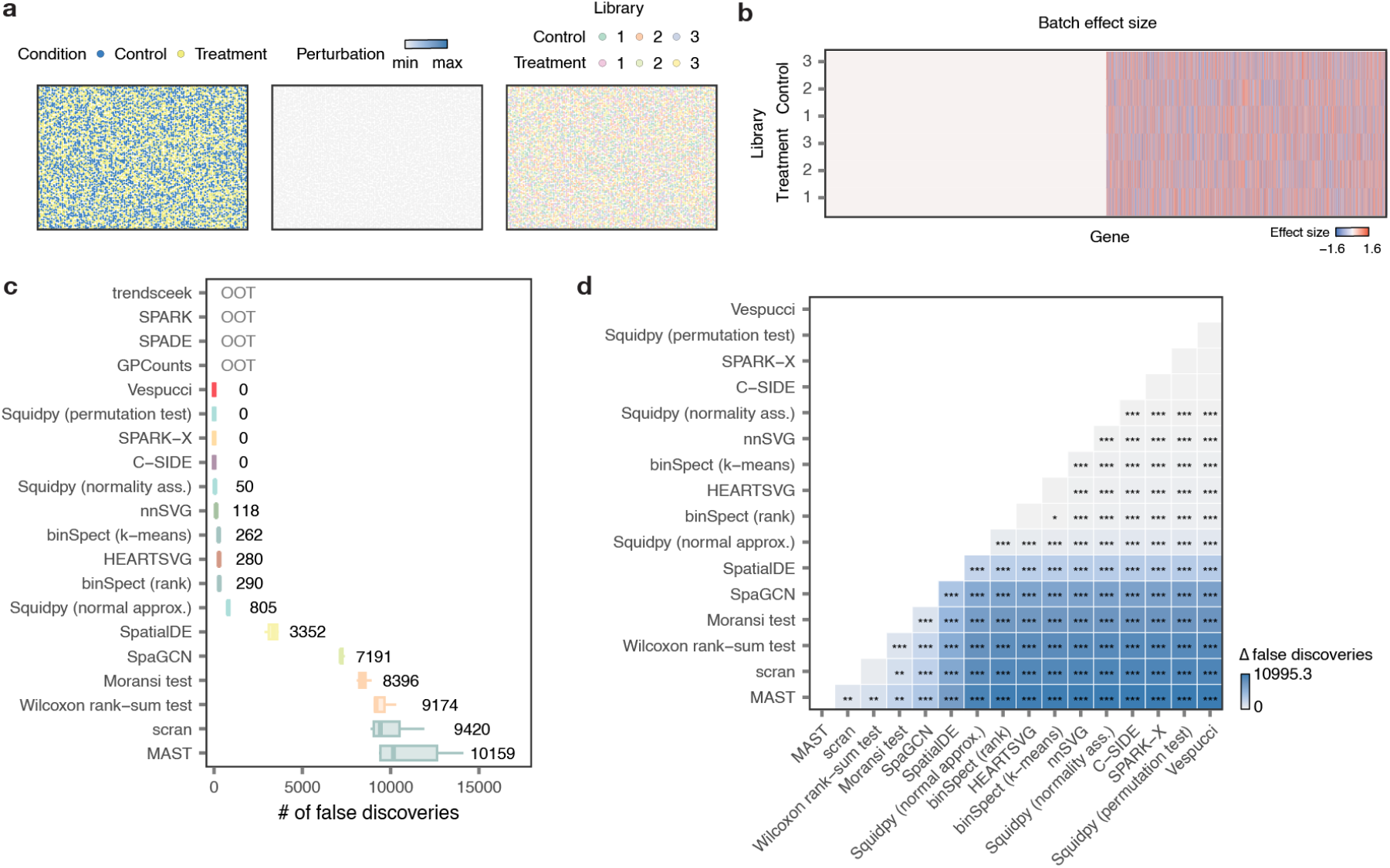
Control of the false discovery rate by computational methods for comparative analysis of spatial atlases. **a-b**, Schematic overview of the simulation framework for testing control of the false discovery rate. Spatial data was simulated without any perturbation effect between two experimental conditions. A total of three libraries per condition were simulated, with 50% of the genes in the dataset affected by batch effects that vary across all six replicates. **c**, Total number of genes identified as DE between conditions by Vespucci versus 19 baseline methods at 5% FDR. **d**, Heatmap comparing the mean number of DE genes for each pair of computational methods. *, p < 0.05; **, p < 0.01; ***, p < 0.001, two-tailed paired t-test.

**Extended Data Fig. 4.**
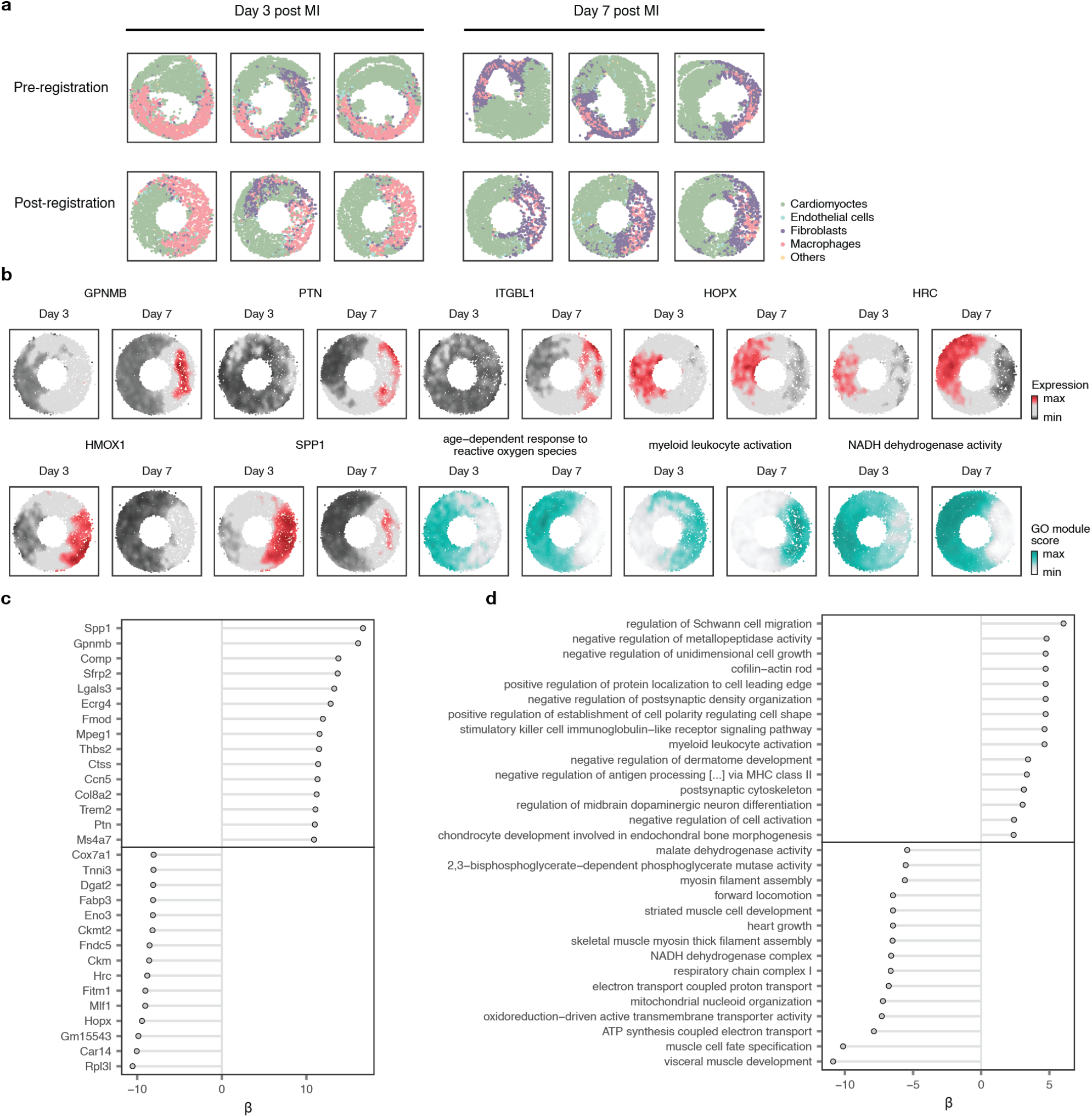
Identification of perturbation-responsive genes and gene programs in myocardial infarction. **a**, Cell type deconvolution of the myocardial infarction dataset using matched single-nucleus RNA-seq from the same mouse model, showing the major cell types of the myocardium. Top, spatial barcodes at their original coordinates on the tissue section; bottom, spatial barcodes after registration to an annular common coordinate framework. **b**, Additional genes and gene programs (Gene Ontology terms) prioritized by Vespucci in the myocardial infarction dataset. **c**, Top-15 genes positively and negatively associated with the AUC assigned by Vespucci at each spatial coordinate. **d**, As in **c**, but for gene programs (Gene Ontology terms).

**Extended Data Fig. 5.**
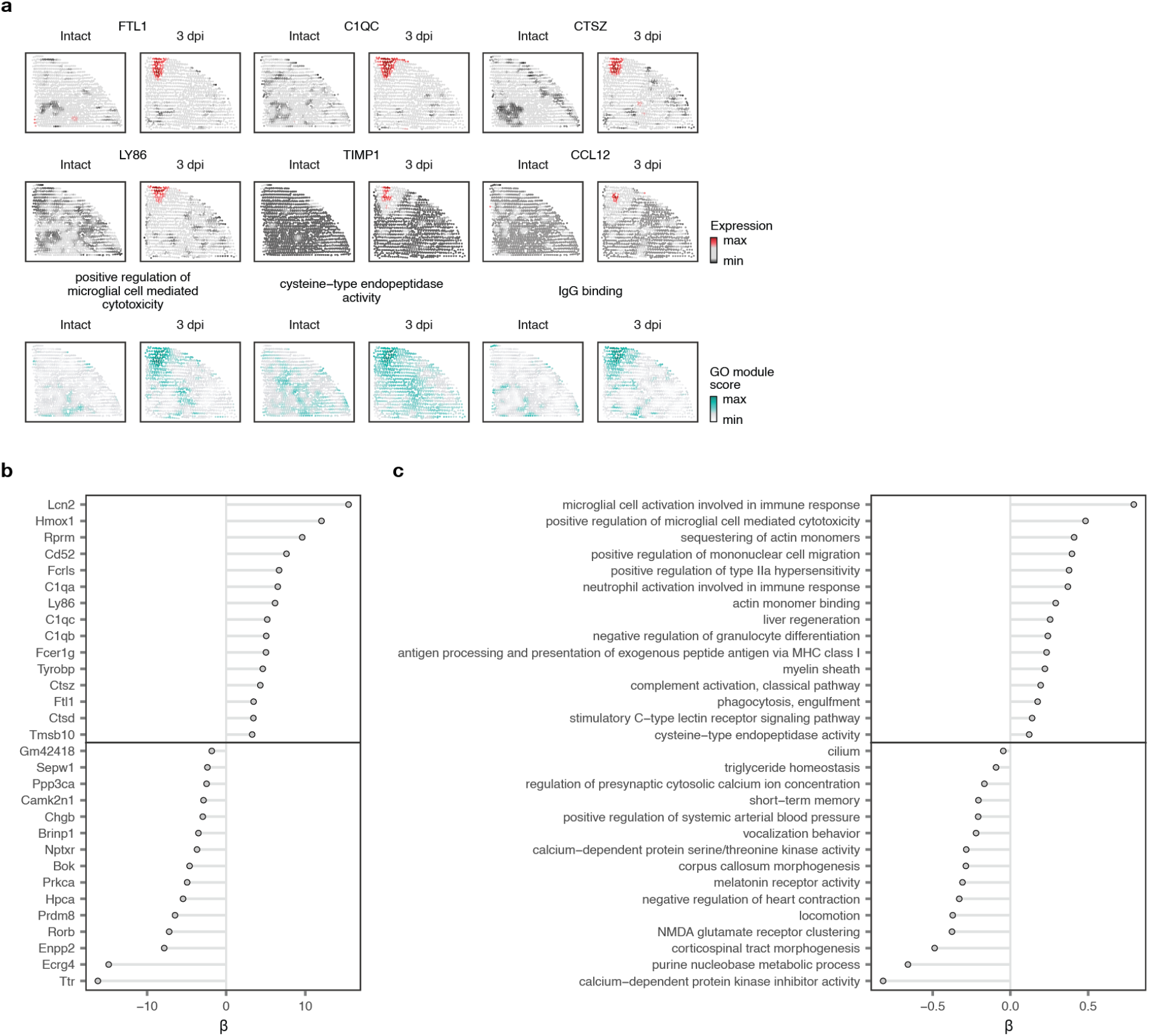
Identification of perturbation-responsive genes and gene programs in traumatic brain injury. **a**, Additional genes and gene programs (Gene Ontology terms) prioritized by Vespucci in the traumatic brain injury dataset. **b**, Top-15 genes positively and negatively associated with the AUC assigned by Vespucci at each spatial coordinate. **c**, As in **b**, but for gene programs (Gene Ontology terms).

**Extended Data Fig. 6.**
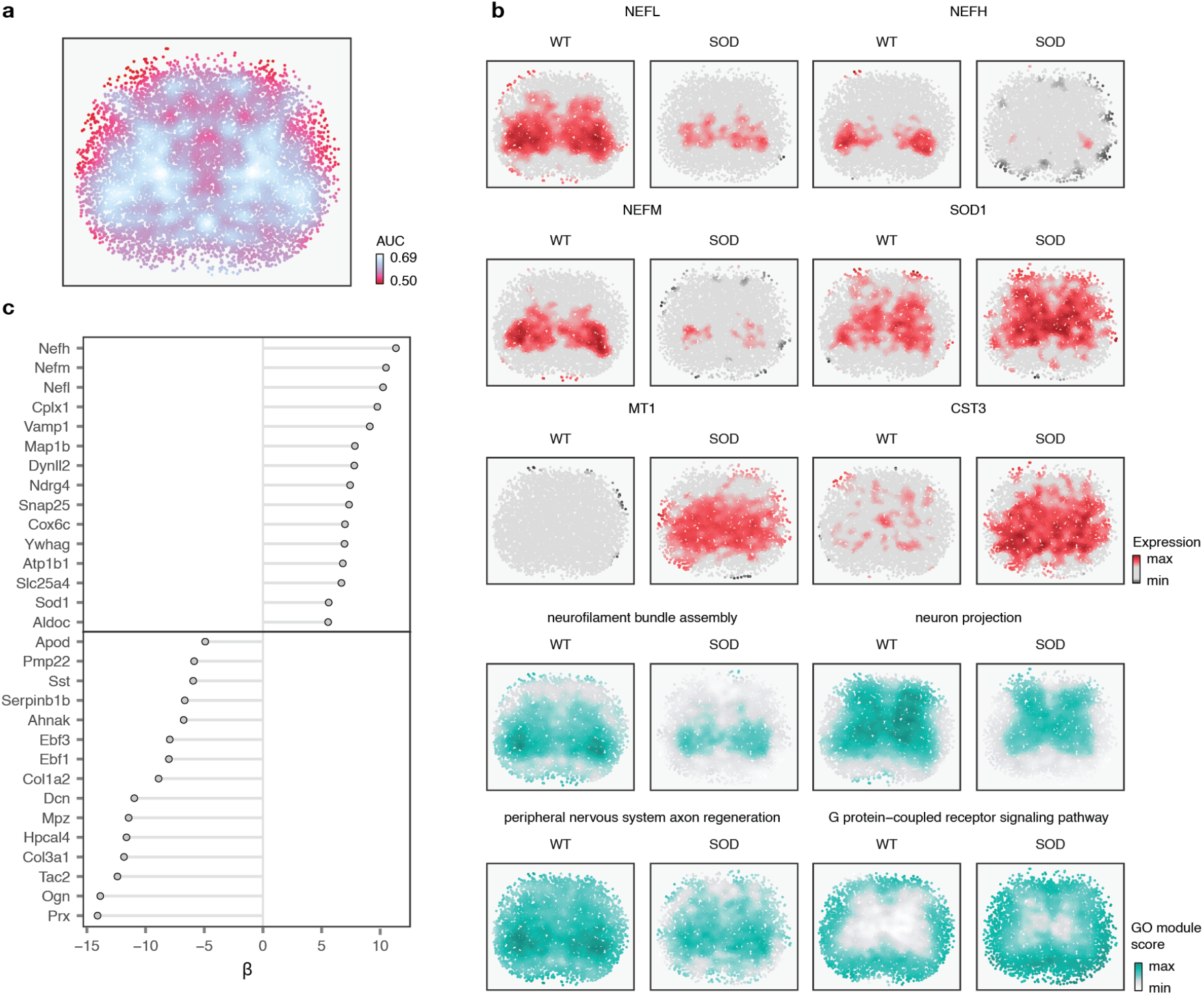
Identification of perturbation-responsive genes and gene programs in ALS. **a**, Spatial prioritization of transcriptionally perturbed regions in the ALS dataset, as quantified by the AUC assigned at each spatial coordinate. **b**, Additional genes and gene programs (Gene Ontology terms) prioritized by Vespucci in the ALS dataset. **c**, Top-15 genes positively and negatively associated with the AUC assigned by Vespucci at each spatial coordinate. **d**, As in **c**, but for gene programs (Gene Ontology terms).

**Extended Data Fig. 7.**
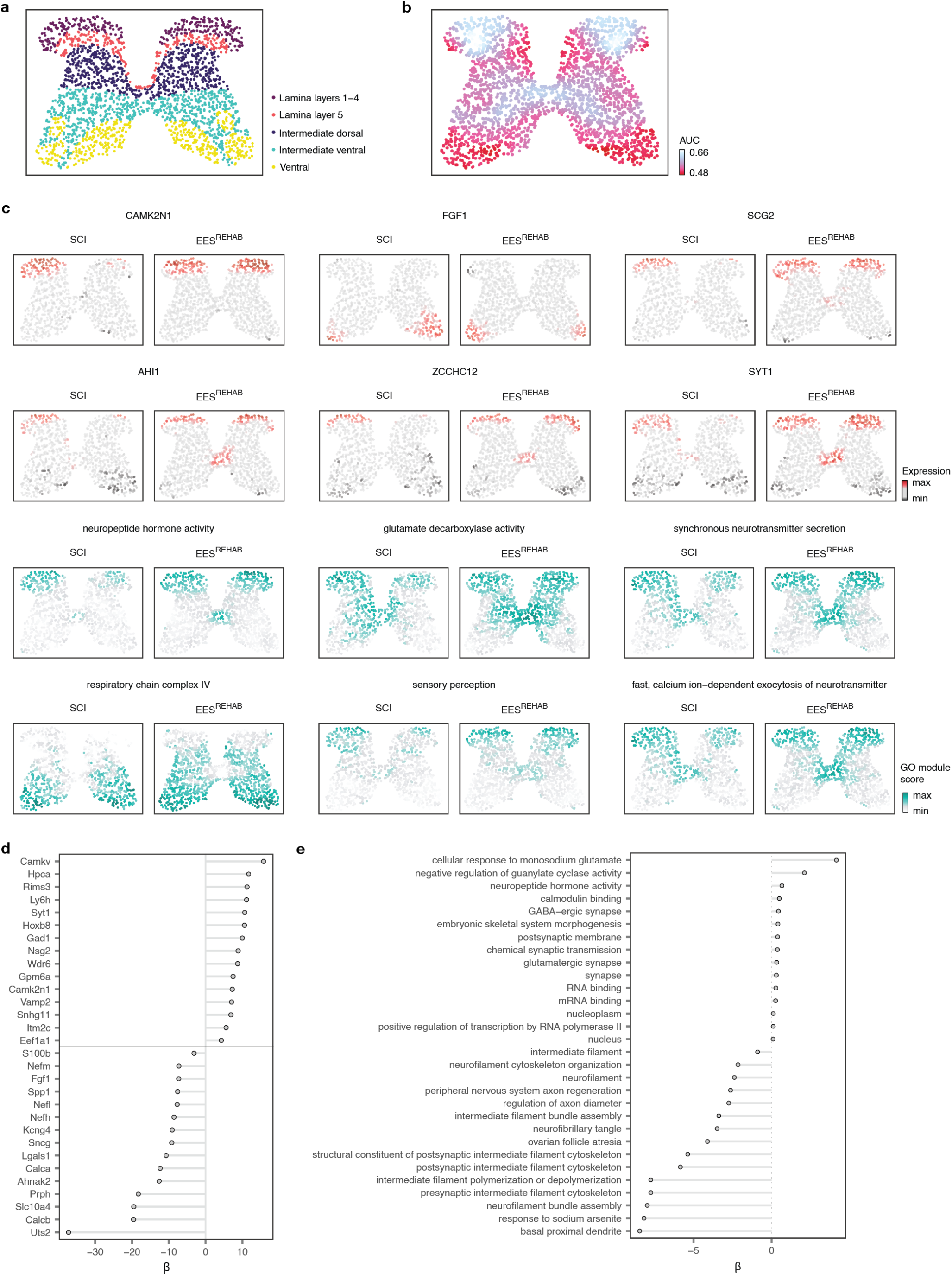
Identification of perturbation responsive genes and gene programs involved in the recovery of walking following neuroprosthetic rehabilitation after spinal cord injury. **a**, Alignment of the neuroprosthetic rehabilitation dataset to the common coordinate system of the spinal cord from the Allen Brain Atlas and the corresponding spinal cord regions assigned to each spatial barcode. **b**, Spatial prioritization of transcriptionally perturbed regions in the neuroprosthetic rehabilitation dataset, as quantified by the AUC assigned at each spatial coordinate. **c**, Selected genes and gene programs (Gene Ontology terms) prioritized by Vespucci in the neuroprosthetic rehabilitation dataset. **d**, Selected genes and gene programs (Gene Ontology terms) prioritized by Vespucci in the neuroprosthetic rehabilitation dataset. **e**, As in **d**, but for gene programs (Gene Ontology terms).

**Extended Data Fig. 8.**
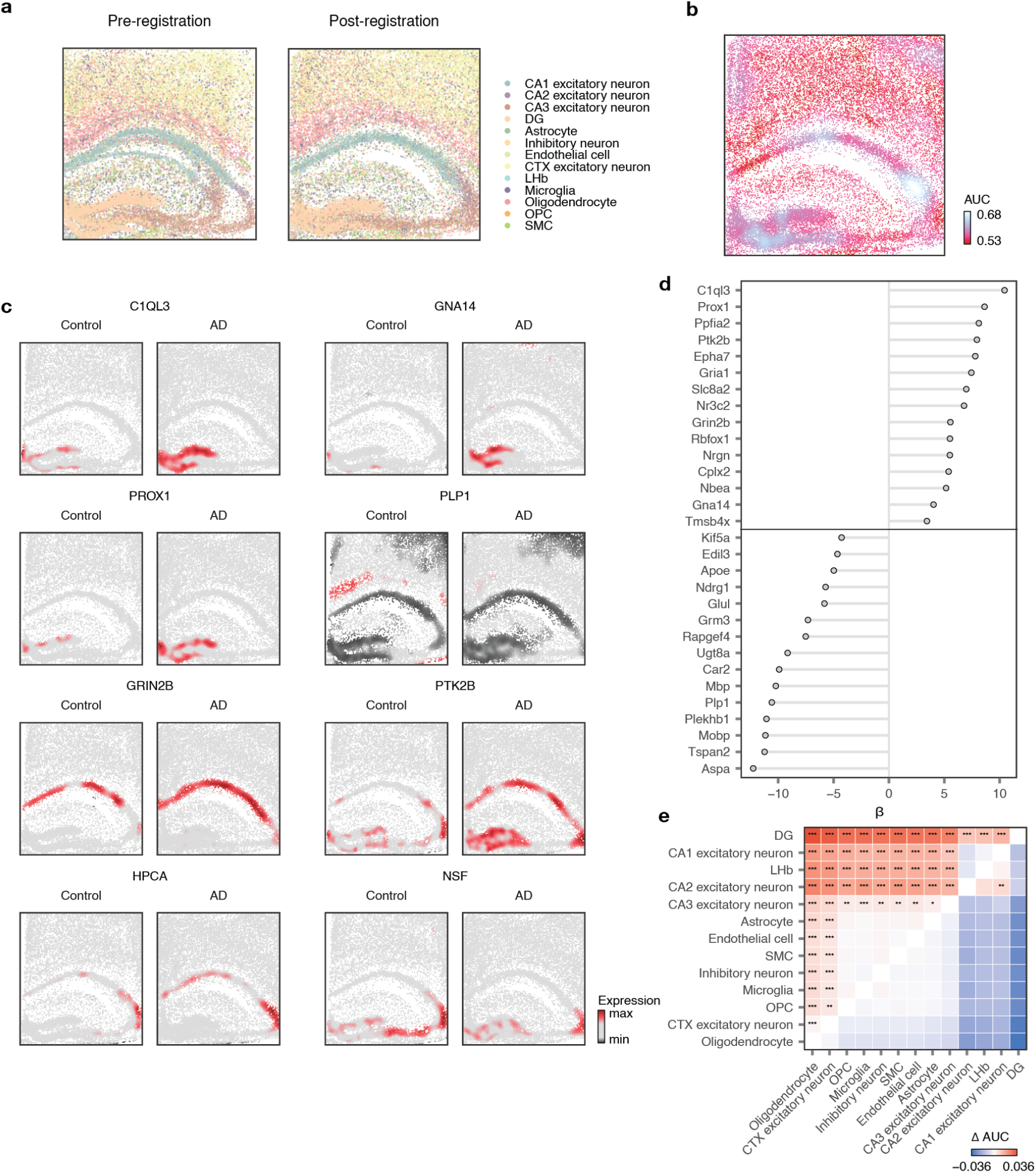
Identification of perturbation-responsive genes and gene programs in Alzheimer’s disease. **a**, Registration of spatial transcriptomics data from the hippocampus of a mouse model of Alzheimer’s disease to a common coordinate framework. **b**, Spatial prioritization of transcriptionally perturbed regions in the Alzheimer’s disease dataset, as quantified by the AUC assigned at each spatial coordinate. **c**, Selected genes prioritized by Vespucci in the Alzheimer’s disease dataset. **d**, Top-15 genes positively and negatively associated with the AUC assigned by Vespucci at each spatial coordinate. **e**, Heatmap comparing the median AUC assigned to each pair of cell types in the Alzheimer’s disease dataset. Vespucci prioritizes cells annotated as dentate gyrus neurons. *, p < 0.05; **, p < 0.01; ***, p < 0.001, two-tailed paired t-test.

**Extended Data Fig. 9.**
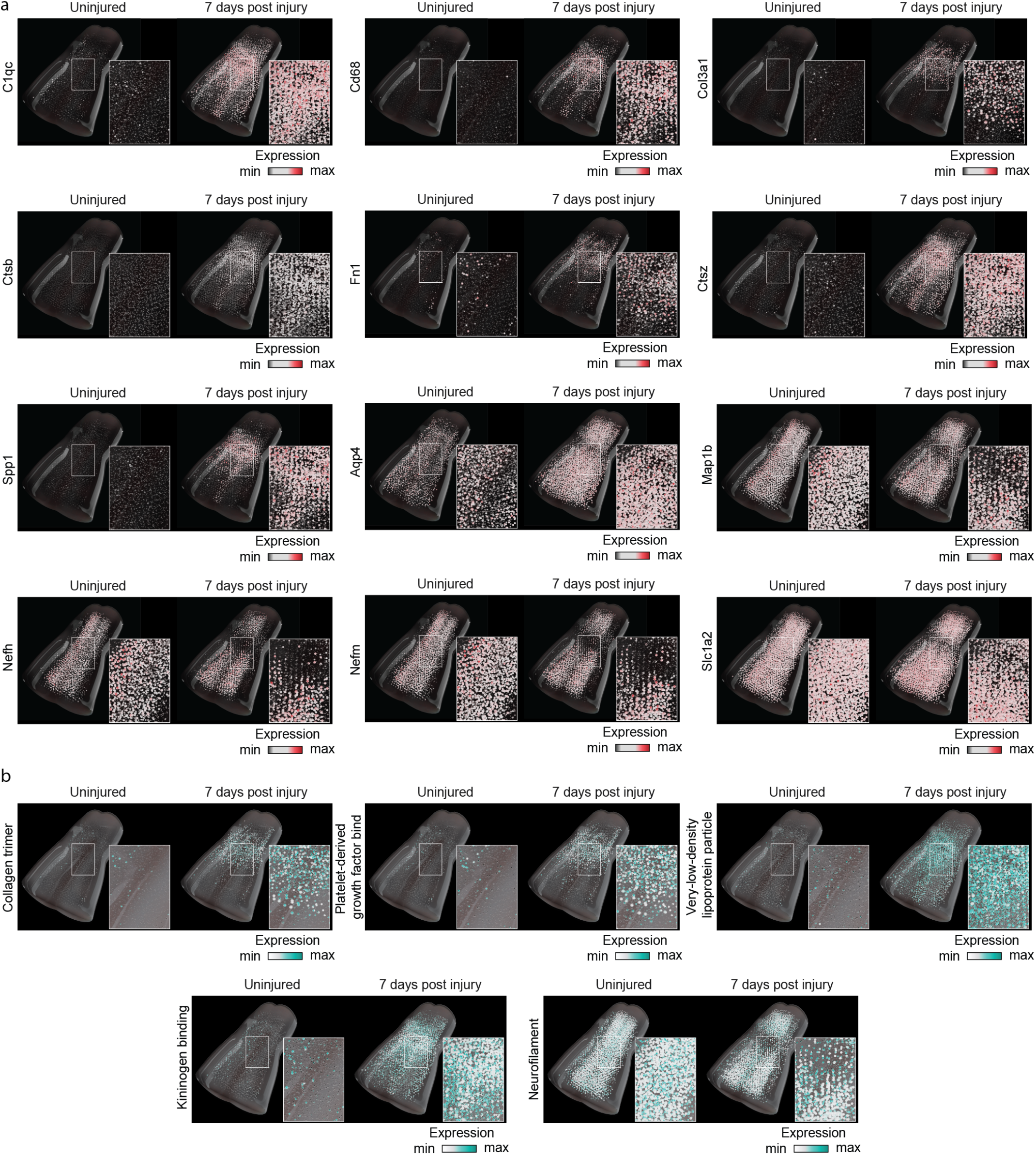
Identification of perturbation-responsive genes and gene programs in a three-dimensional comparative spatial atlas. **a**, Selected genes prioritized by Vespucci in the 3D spinal cord injury dataset. **b**, Selected gene programs (Gene Ontology terms) prioritized by Vespucci in the 3D spinal cord injury dataset.

**Extended Data Fig. 10.**
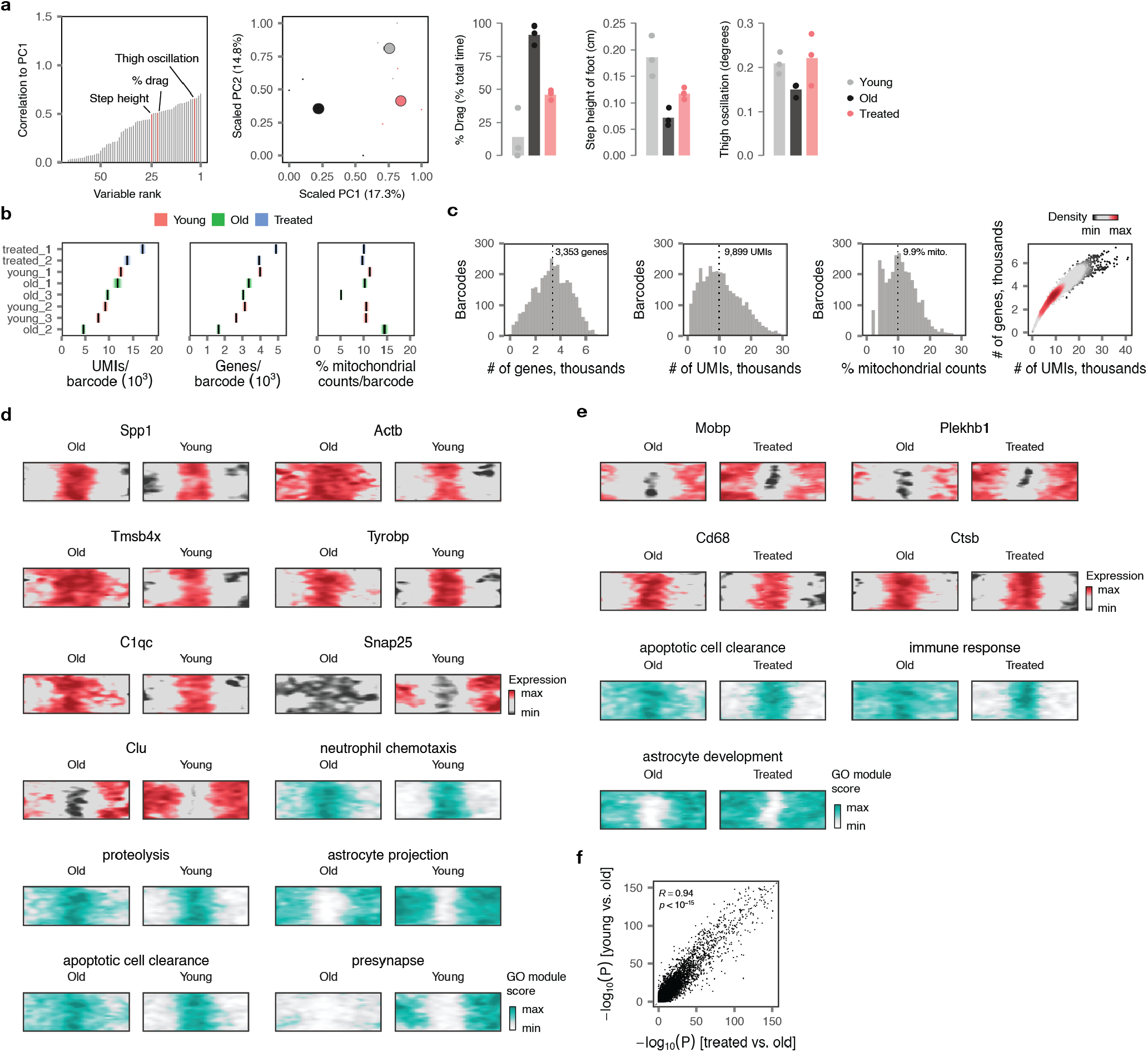
A comparative spatial transcriptomics atlas of the injured spinal cord in young and old mice. **a**, Walking performance in young mice, and old mice with and without treatment. Walking performance was quantified using principal component analysis applied to 55 gait parameters calculated from kinematic recordings. Small points show individual gait cycles (n > 10 per mouse, n = 3 mice per group). Large points show the mean of each experimental group. The first principal component (PC1) distinguished gaits from mice across different experimental groups. Analysis of factor loadings on PC1 revealed that the percentage of paw dragging, step height, and the extent of thigh oscillation, were among the parameters that showed high correlations with PC1. Bars show the mean values of each gait parameter per group. **b**, Section-level quality control statistics for spinal cord sections profiled by spatial transcriptomics. Left, mean number of UMIs per barcode; middle, mean number of genes detected per barcode; right, mean proportion of mitochondrial counts per barcode. Lines and shaded areas show the mean and standard deviation, respectively, per section. **c**, Barcode-level distributions of quality control statistics over all sections (far left, number of UMIs per barcode; middle left, number of genes detected per barcode; middle right, proportion of mitochondrial counts per barcode; far right, relationship between the number of UMIs and number of genes detected per spatial barcode). **d**, Additional genes and gene programs (GO terms) prioritized by Vespucci in the comparison of spinal cords from young versus old injured mice. **e**, Additional genes and gene programs (GO terms) prioritized by Vespucci in the comparison of spinal cords from old versus treated injured mice. **f**, Correlation between –log_10_ p-values assigned to each gene by Vespucci in comparisons of young versus old injured mice, y-axis, as compared to old versus treated injured mice, x-axis.

## Supplementary Algorithm 1: Vespucci algorithm

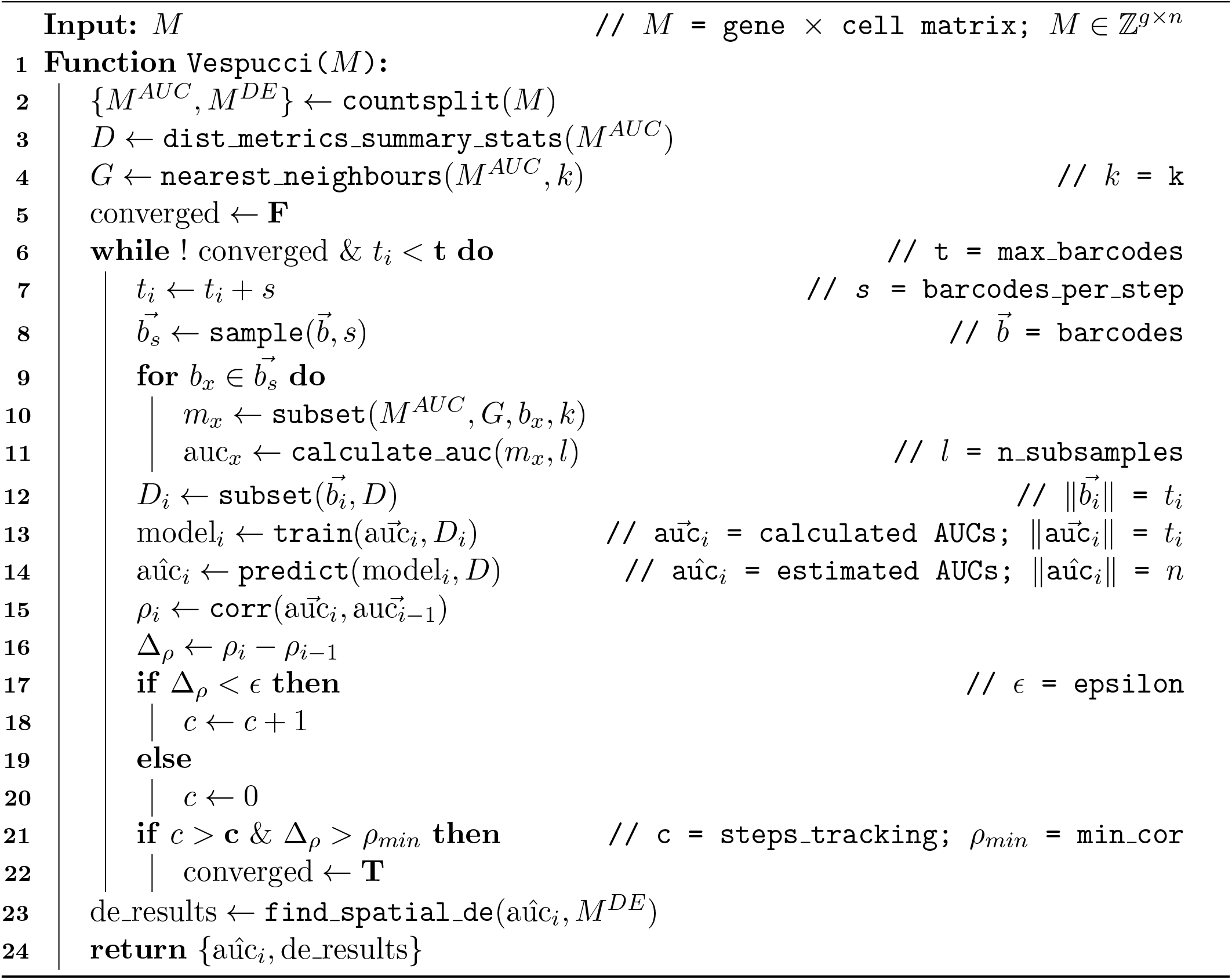

